# Emergence of bluetongue virus serotype 3 in the Netherlands in September 2023

**DOI:** 10.1101/2023.09.29.560138

**Authors:** Melle Holwerda, Inge M.G.A. Santman-Berends, Frank Harders, Marc Engelsma, Rianka P.M. Vloet, Eveline Dijkstra, Rene G.P. van Gennip, Maria H. Mars, Marcel Spierenburg, Lotte Roos, René van den Brom, Piet A. van Rijn

## Abstract

Since 1998, WOAH-notifiable bluetongue virus (BTV) serotype 1, 2, 3, 4, 6, 8, 9, 11, and 16 have been reported in Europe. In mid-August, 2006, a BTV-8 outbreak started in Northwestern Europe. The Netherlands was declared BT-free again in February 2012 and annual monitoring demonstrated BT-freedom up to 2023. On September 3^rd^ 2023, clinical manifestations in sheep typical for BT were notified to the Dutch Food and Product Safety Consumer Authority. Laboratory diagnosis confirmed BTV-infection on September 6 and the first notifications of clinical signs in cattle were also reported. Two days later, the virus was identified as serotype 3 by whole genome sequencing. Clinical signs were like these of BTV-8 outbreak and were most serious in sheep. Retrospective analysis revealed no earlier circulation of BTV. It was concluded that the BTV-3 outbreak was detected shortly after introduction, while the virus source and route of introduction remains unknown.

**Article summary line:** During September 2023, a novel serotype 3 Bluetongue virus (BTV) emerged in the Netherlands which affected sheep and cattle.

## Introduction

Bluetongue virus (BTV) is an arthropod-borne virus, which can cause clinical disease and mortality in ruminants. Exclusively, all types of ruminants are susceptible for infection with BTV, while infections in new world camelids have incidentally also been described (1–3). Other species including humans are not susceptible for infection, indicating that BTV is not a zoonosis.

BTV is transmitted by certain species of *Culicoides* midges and was historically only present between latitudes 35°S and 50°N (4,5). The BTV serogroup consists of more than thirty serotypes, which show no or limited cross-protection between serotypes. Since 1998, several BTV serotypes (1, 2, 3, 4, 6, 8, 9, 11 and 16) have been present in Europe and the Mediterranean Basin (6). In 2006, bluetongue virus serotype 8 (BTV-8) emerged in northwestern Europe for the first time, and the Netherlands was the first country where the infection was detected (7,8). After a major BTV-8 outbreak in 2006 and 2007, in 2008 an emergency BTV-8 vaccine became available (9,10) and many cattle herds and small ruminant flocks participated in the voluntary vaccination program that was implemented by the Dutch government (11). This resulted in a dramatic decline in the number of clinical notifications at the Dutch Food and Consumer Product Safety Authority (NVWA) in 2008. At the end of 2008, over 80% of the susceptible host population tested positive for antibodies due to natural infection or vaccination. From 2009 on, no new infections were observed and after three years of screening of possible BTV circulation, the Netherlands regained its official BT-free status in February 2012. Since then, this disease-free status was monitored annually according to EU regulation 1108/2008/EC and was confirmed without interruption up to December 2022. However, based on the risk of introduction of BTV-8 from neighboring countries, vaccination was allowed and therefore some farmers still vaccinated their animals for serotype 8.

On 3 September 2023, clinical signs in sheep indicative for BT were notified to the authorities by two veterinary practices located at the middle of the Netherlands at the same time. This paper describes the actions that were taken after the first notified clinical case was confirmed as infection with BTV.

## Methods

### Sheep and cattle population in the Netherlands and clinical examination

In 2022, around 1,080,631 sheep and 1,596,894 dairy cattle (>2 year)were present at in approximately 31,000 sheep farms and 14,000 cattle herds in the Netherlands (12,13). Farms with a suspicion of BT that notified to the authorities were visited by a veterinary team with specialists in (small) ruminant health to review reported clinical symptoms and to take samples for BTV diagnostics. Additionally, several farms were visited by GD that were already confirmed as BTV-positive, and sheep and cattle on these farms were clinically examined and the clinical signs were described.

### Realtime PCR

The real-time PCR was performed according to the protocol of van Rijn *et al*. (2012) (14). Briefly, viral RNA was extracted from 200 μl of EDTA-blood using the Magnapure 96 robotic machine (Roche, Basel, Switzerland) in combination with the MagnaPure 96 DNA and viral NA small volume kit (Roche). For RT-PCR, five microliter of elution was loaded into a 96-well plate with the Light Cycler RNA master hybridization Probe kit (Roche) and amplification was performed in a Lightcycler 480 machine (Roche) using integrated software version 1.5.1.

### Competition ELISA on individual samples

The competition ELISA was performed with the ID screen bluetongue competition enzyme-linked immunosorbent assay (ELISA) according to the manufacturers protocol (Innovative Diagnostics, Montpellier, France). Optical density was measured at 450 nanometer using a Multiskan FC machine (Thermo Scientific, USA) in combination with Mikrowin (software version 5.09) and % blocking was calculated using the positive and negative control supplied with the kit.

### SISPA-based whole genome Sequencing using Oxford Nanopore Technology

RNA extracted from EDTA-blood was applied to amplify the viral genomic segments using a Sequence Independent Single Primer Amplification (SISPA) approach. Initial first strand cDNA synthesis was performed with five μl of RNA in combination with Superscript III (Thermo Fischer) according to manufacturer’s protocol using 2 uM of 5’-GTT TCC CAG TCA CGA TA N9-3’. The mixture was incubated for 3 minutes at 95 ºC to denaturate viral double-stranded RNA followed by cooling on ice. The remaining ingredients were added to the reaction and incubated at 25 ºC for 5 minutes, 42 ºC for 50 minutes, 70 ºC for 15 minutes and stored at 4 ºC. Second strand synthesis was performed using Sequenase (Thermo Fischer) according to manufacturer’s protocol. Amplification of the products was performed using Q5 high-fidelity DNA polymerase (New England Biolabs, USA) according to the guidelines of the producer with 2 μM of 5’-GTT TCC CAG TCA CGA TA-3’. The following cycle conditions were used: 94 ºC for 4 minutes, 68 ºC for 5 minutes following 35 cycles of 94 ºC for 30 seconds, 50 ºC for 1 minute, 68 ºC for 3 minutes, followed by 68 ºC for 5 minutes and cooldown to 10 ºC. To enhance the number of viral reads, a size selection of >200 base pairs was performed using the SPRIselected beads (Beckman-Coulter, USA) with a ratio of 0.8. From each individual sample was ∼150 nanogram barcoded using the Native barcoding v14-kit (SQK-NBD114.96, Oxford Nanopore technologies, United Kingdom) according to manufacturer’s protocol. Nanopore sequencing was performed on an Oxford Nanopore Promethion Flow Cell R10 (M version). To align the reads, Minimap 2(V2.26) was applied against a custom BTV reference database to construct a draft genome using reference-based mapping (15). The sequences were deposited on NCBI on 26 September 2023 with accession numbers OR603992-OR604001.

Phylogenetic analysis of the obtained genome sequences was performed for each genome segment separately with the top-15 BLAST (NCBI GenBank accession date September 11, 2023) results included in the analysis. In addition, Seg-2 reference strains and a selected number of closely related BTV-3 strains were added in the phylogenetic analysis (16). The sequences were aligned using MAFFT v7.475 (17) followed by reconstructing the phylogeny using maximum likelihood (ML) analysis with IQ-TREE software v2.0.3 (18) and 1,000 ultrafast bootstrap replicates (19), and visualizing the ML tree using the R package ggtree (20).

### Bulk tank milk ELISA

For the retrospective analysis of BTV-antibodies in bulk milk, the indirect ELISA based on the recombinant VP7 protein (ID Screen ® Bluetongue Milk Indirect, Innovative Diagnostics, Montpellier, France) was used according to manufacturer’s protocol. This ELISA has been validated in 2007 in the Netherlands (21). To study the herd prevalence of Bluetongue serotype 3 infections in 2023, the cut off was used as prescribed in the manual: S/P≤30% is considered negative, 30>S/P%≤40% is considered doubtful and S/P>40% is considered positive.

### Retrospective study

To investigate whether the initial outbreak started in the area of the four BTV-3 confirmed sheep farms in the middle of the Netherlands, bulk tank milk samples from cattle from all over the Netherlands that were submitted for routine testing in August were screened for the presence of BTV-antibodies. Royal GD coordinates a national monitoring program for which approximately 90% (N=12,000) of the Dutch dairy cattle farmers submit bulk tank milk samples on a monthly base. To gain insight into the presence of BTV-antibodies in Dutch dairy herds and thereby indicating an earlier infection then September 2023, thousand milk-samples from all over the Netherlands were tested.

Identification and registration data was used (RVO, Assen the Netherlands) to enable selection of dairy herds that did not purchase any cattle during the vector active season in 2023 (from April on) and that only housed Dutch bred animals (N=7,900). The Netherlands is divided into twenty compartments as proposed in Commission Decision 2005/393/EC and the 1,000 bulk milk samples were randomly selected stratified to the twenty compartments. This resulted in approximately fifty sampled herds which enabled a prevalence estimate with a precision of 14% and 95% confidence. On 11 September the first preliminary results were presented to the government. On 13 September additional data on vaccination purchases that are registered in the MediRund database (ZuivelNL, the Hague the Netherlands) became available and were combined with the results of the bulk milk screening.

### Map generation

The sheep and cattle density are graphically displayed in thematic maps of the Netherlands, with the density presented at two-digit postal code level. BTV-confirmed clinical notifications of sheep and cattle cases are presented as dots in the respective maps until 29/09/2023. All maps are generated using Stata version 17^®^ (Statacorp, 2021).

## Results

### Timeline **(Figure 1)**

On 3 and 4 September 2023, NVWA was notified of clinical signs that were indicative for BTV infection at five sheep farms in the middle of the Netherlands near the “Loosdrechtse plassen”. Flocks were visited by a team of veterinary specialists, where serum and EDTA-blood were collected from sheep which were send to the Dutch National Veterinary Reference Laboratory for BTV, Wageningen Bioveterinary Research (WBVR).

**Figure 1.**
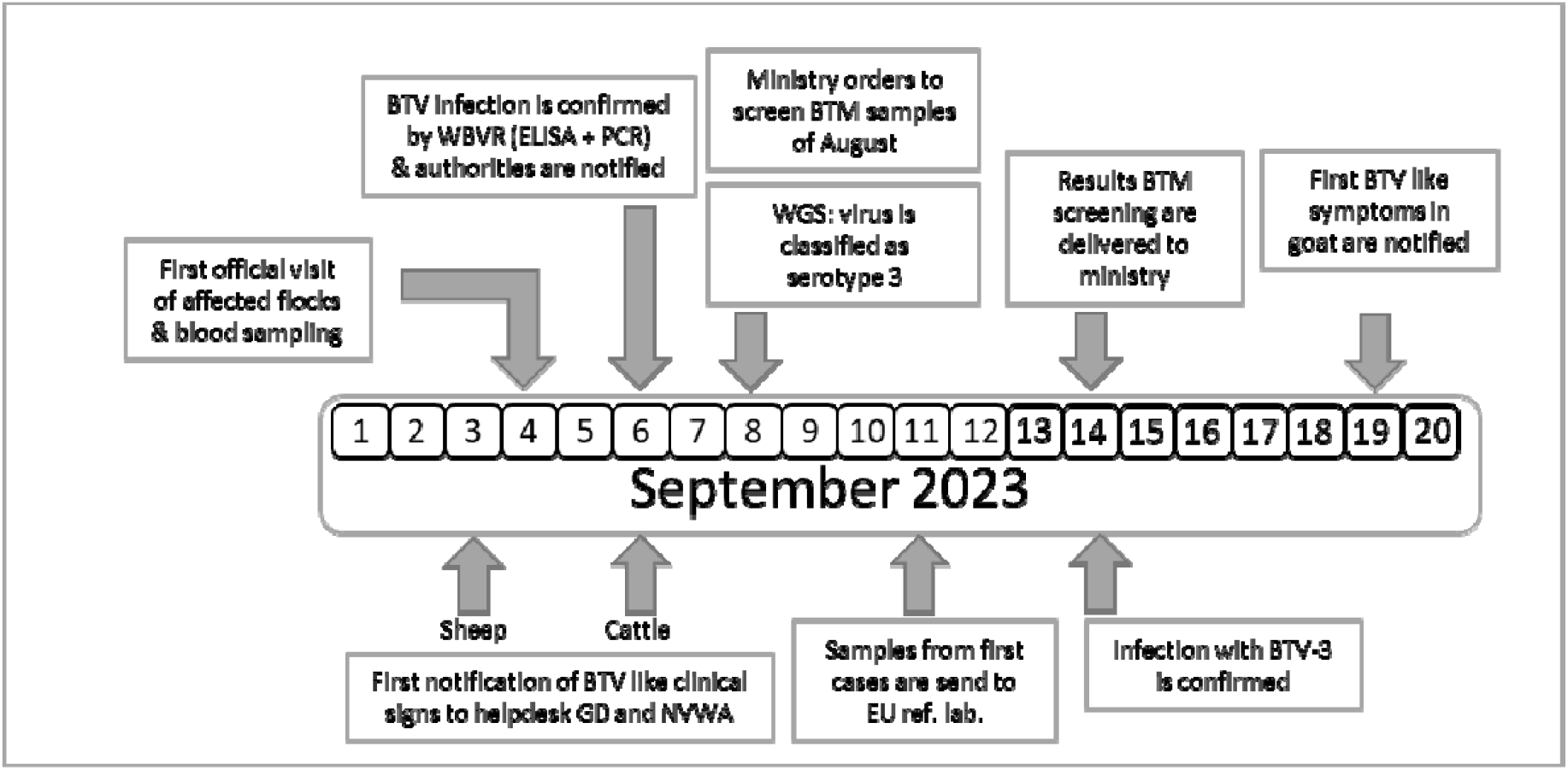
Graphical representation of the timeline of the initial BTV-3 outbreak in the Netherlands in September 2023. Abbreviations: BTM: bulk tank milk, BTV: Bluetongue virus, WGS: Whole genome Sequencing.

On 6 September, BTV infections was confirmed by real-time PCR and competition ELISA. Of the seven blood samples taken from five sheep farms, six samples from four different farms tested PCR-positive with Ct-values ranging from 23 to 31. Five out of six PCR-positive sheep also tested positive for antibodies against BT with blocking percentages >90%. Differential diagnosis for BTV include foot-and-mouth disease (FMDV), therefore, the samples were also tested with an inhouse developed and validated PCR-test for FMDV, but no Ct-values were measured. Results were immediately reported to the Dutch Ministry of Agriculture, Nature, and Food quality. Additionally, new samples were requested for confirmation and for shipment to the European Reference Laboratory for BTV, Animal Health Research Center CISA-INIA in Madrid, Spain. In addition, whole genome sequencing was initiated on the PCR-positive samples using Oxford Nanopore Technology by WBVR.

The first suspicion of BTV in cattle was notified to the NVWA on September 6.

On 8 September, three blood samples of sheep sampled at three unrelated farms showed sufficient coverage per nucleotide, ranging from 30 to 2570, (**Table 1**) to reliably determine contig sequences of all ten genome segments for three samples. Contig sequences derived from individual samples were 100% identical. Contigs represented full length sequences of genome segment 1-9 (Seg-1 to Seg-9), including the 5’ and 3’-termini. Contigs of Seg-10 were incomplete and additionally completed by sanger sequencing, except for the ultimate 22 nucleotides at the 3’-end corresponding to the amplification primer. Full length sequences of Seg-1 to Seg-10 are submitted to the NCBI Genbank accession numbers OR603992-OR604001. Phylogenetic analysis of Seg-2 encoding the serotype dominant VP2-protein with the prototypic isolates of notifiable BTV serotypes 1-24 identified the causative agent as BTV serotype 3. As can be seen in **Figure 2A**, phylogenetic clustering was also observed with serotype 13 and 16, confirming previous genetic analysis (22). Based on the phylogenetic tree, WBVR announced that genotyping revealed that the BTV outbreak was caused by serotype 3, due to the high homology with known serotype 3 isolates. Detailed phylogenetic analysis showed a close relationship with Seg-2 from recent Italian and Tunisian BTV-3 isolates **(Figure 2B)**.

**Table 1:**
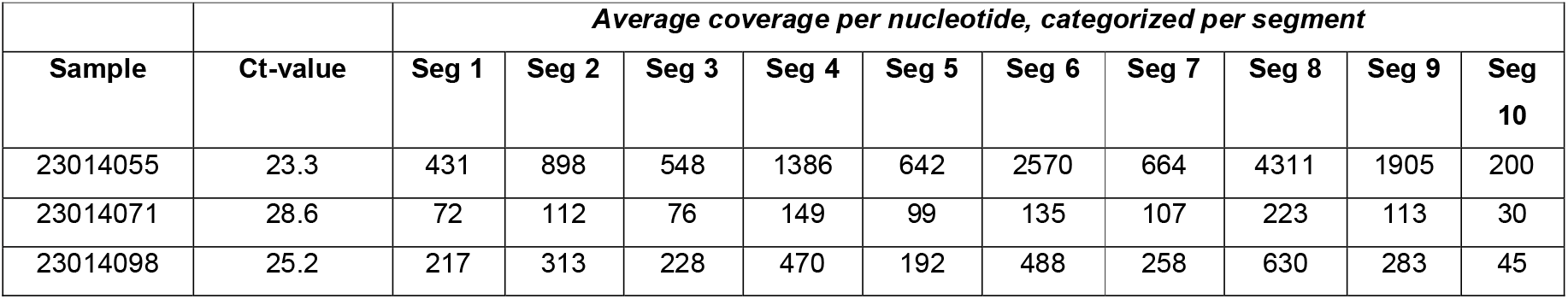
The average coverage of each specific segment of the three samples that were subjected to whole genome sequencing.

| Sample | Ct-value | Average coverage per nucleotide, categorized per segment |  |  |  |  |  |  |  |  |  |
| --- | --- | --- | --- | --- | --- | --- | --- | --- | --- | --- | --- |
|  |  | Seg 1 | Seg 2 | Seg 3 | Seg 4 | Seg 5 | Seg 6 | Seg 7 | Seg 8 | Seg 9 | Seg 10 |
| 23014055 | 23.3 | 431 | 898 | 548 | 1386 | 642 | 2570 | 664 | 4311 | 1905 | 200 |
| 23014071 | 28.6 | 72 | 112 | 76 | 149 | 99 | 135 | 107 | 223 | 113 | 30 |
| 23014098 | 25.2 | 217 | 313 | 228 | 470 | 192 | 488 | 258 | 630 | 283 | 45 |

**Figure 2:**
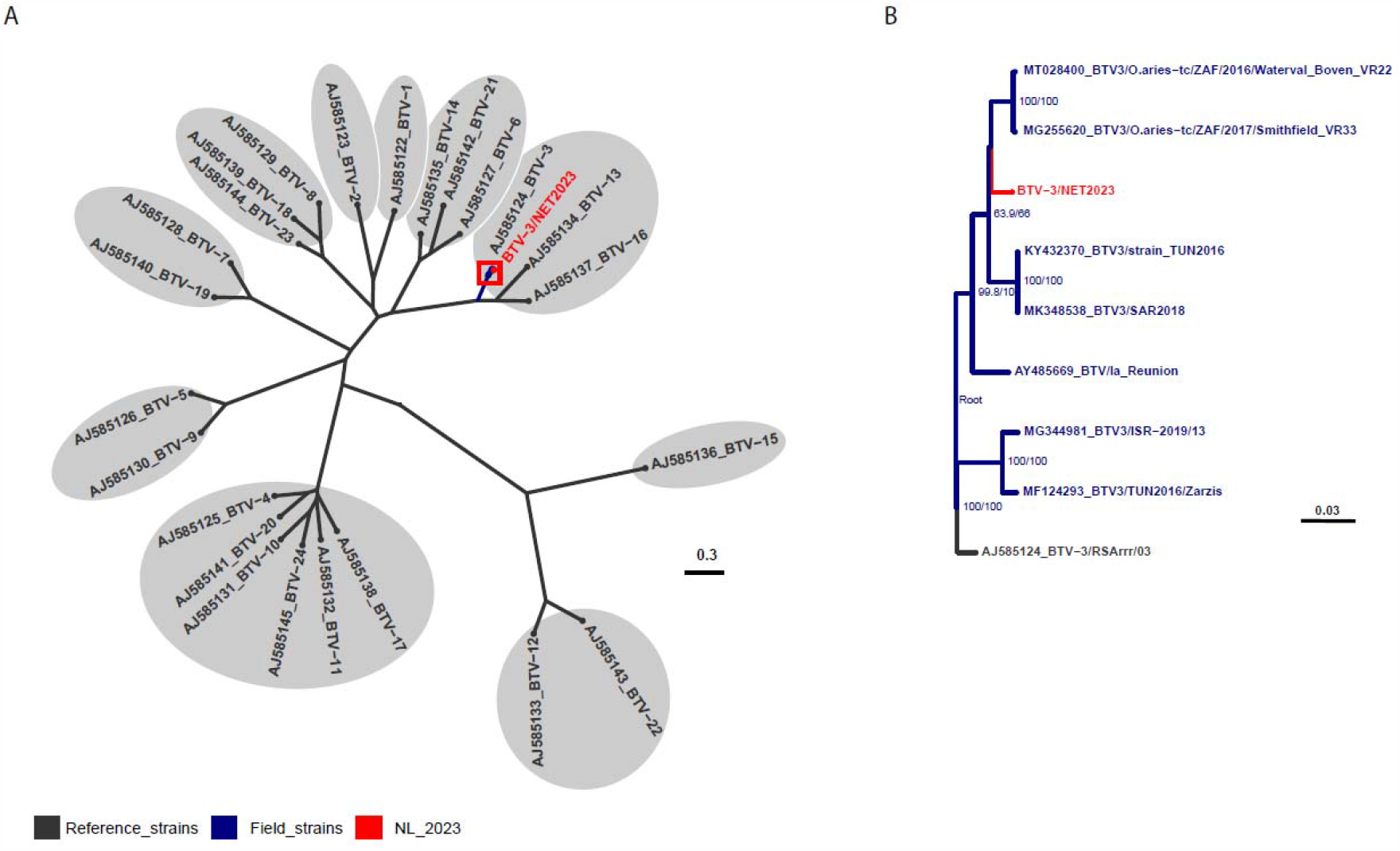
Phylogenetic trees obtained with the Maximum Likelihood method showing (A) the obtained segment 2 sequence with reference strains from each of the 24 notifiable BTV serotypes and in detail (B) selected field strains of closely related BTV-3 sequences. Unrooted trees with UFBoot2 bootstrap values indicated at the nodes, GenBank accession numbers included in sequence name.

Phylogenetic analyses for other individual genome segments of BTV-3/NET2023 did not show a clue of a particular ancestor but the closest identity (>97%) to genome segments of various BTVs **(Table 2)**.

**Table 2:** The percentage of homology between BTV-3/NET2023 with the closest isolate, which is deposited on NCBI.

| Genome segment | Virus protein | Highest % identity with BTV-3/NET2023 | Isolate name | Accession number |
| --- | --- | --- | --- | --- |
| Seg-1 | VP1 | 97,69 | BTV-8/2020_13 | OQ860824.1 |
| Seg-2 | VP2 | 98,09 | BTV-3/ZIM2002/01 | AJ585179.1 |
| Seg-3 | VP3 | 98,3 | BTV-5/O.aries-tc/ZAF/2011/Benoni_01012015 | MG255451.1 |
| Seg-4 | VP4 | 98,37 | BTV-3/TUN2016/Zarzis | MF124295.1 |
| Seg-5 | NS1 | 98,42 | BTV-1/ISR-2050/19 | OM502356.1 |
| Seg-6 | VP5 | 97,5 | BTV-3/O.aries-tc/ZAF/2017/Smithfield_VR33 | MG255623.1 |
| Seg-7 | VP7 | 98,27 | BTV-3/O.aries-tc/ZAF/2016/Waterval_Boven_VR22 | MT028405.1 |
| Seg-8 | NS2 | 97,69 | BTV-2/O.aries-tc/ZAF/2017/Queenstown_VR18 | MG255577.1 |
| Seg-9 | VP6/NS4 | 97,43 | BTV-4/SPA2003/03 | KP821911.1 |
| Seg-10 | NS3/NS3a | 98,42 | BTV-18/BT32/76 | JX272448.1 |

On 11 September, newly collected serum and EDTA-blood samples from the four initial farms were subjected for confirmation and also send towards the EURL for confirmation and serotyping by serotype-specific real-time PCR tests.

On 14 September, the EURL confirmed the results based on the WOAH-recommended PCR-test targeting seg-10. Serotype-specific real-time PCR tests specific for serotype 3, 4 and 8 clearly confirmed serotype 3. This result was immediately forwarded to the Ministry of LNV and NVWA.

On 19 September, samples of the first suspicion of BTV-like signs in a goat was reported. In addition, virus isolation of BTV-3/NET2023 from EDTA-blood derived from sheep from the initial four farms was shown to be successful on KC-cells (23) .

### Clinical manifestation in sheep and cattle

Sheep in flocks that were among the first ones to report clinical signs, showed signs of fever, lethargy, hypersalivation, ulcerations and erosions of the oral and nasal mucosal membranes, facial oedema, and lesions of the coronary band, lameness and mortality **(Figure 3)**. In the days after the initial confirmation of the outbreak, clinical signs were also reported in cattle. Clinical signs observed in cattle consisted of fever, apathy, conjunctivitis, nasal discharge, erosions and crust formation on lips and nostrils, ulcerations and erosions of oral mucosa, oedema of the nose, coronitis and superficial necrosis of teats.

**Figure 3:**
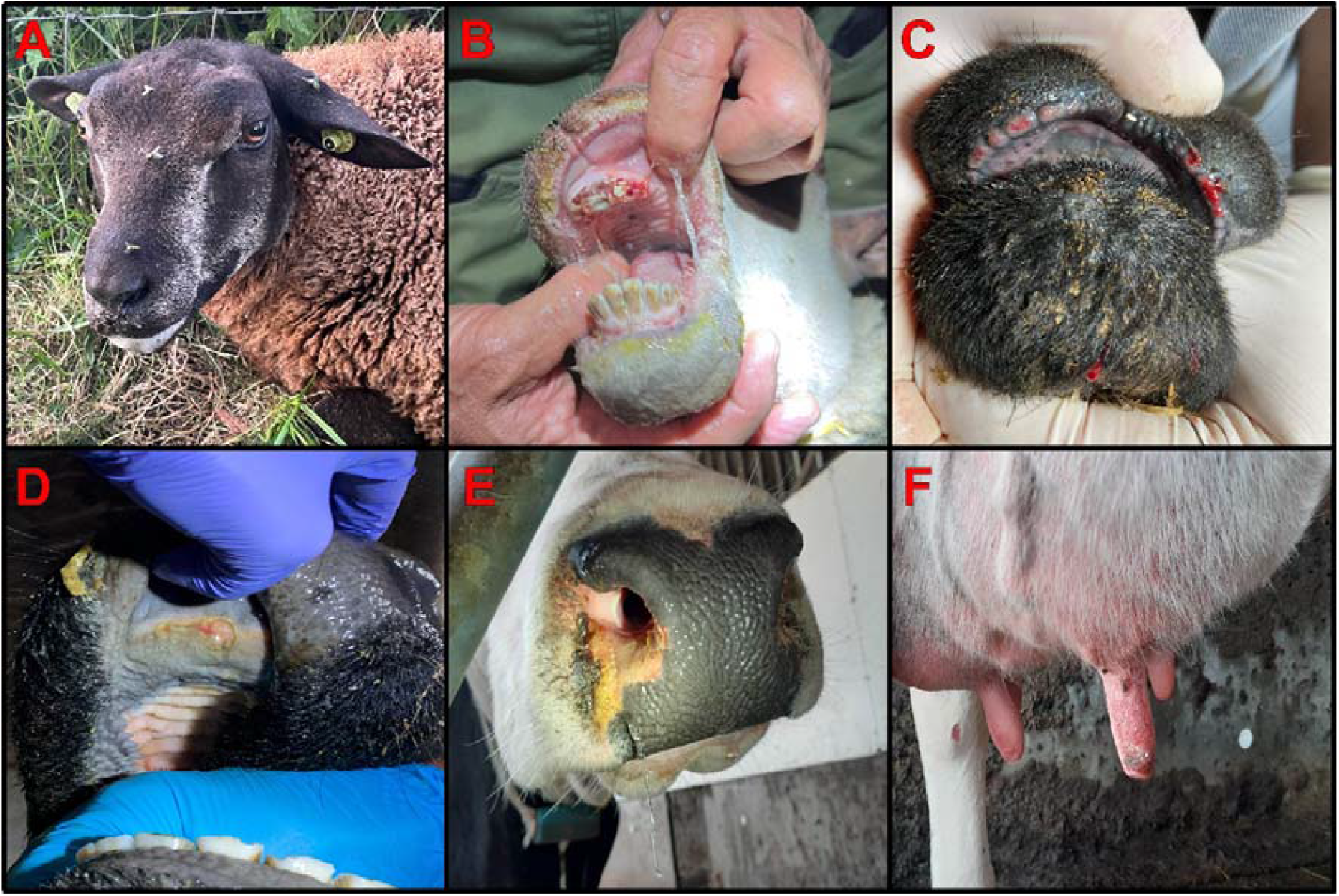
Clinical manifestations of BTV-3/NET2023 in sheep and cattle. (A) Hypersalivation and (B) erosion of the oral mucosal membranes and (C) bleeding of the lips are observed in sheep. In cattle are (D) ulceration on the oral mucosal membrane, (E) crust formation at the nostrils and (F) necrosis of the teats detected.

After the start of the outbreak, the number of notifications increased rapidly in both sheep flocks and cattle herds. The initial cases were four sheep farms. One week later (week 36), in total 25 sheep and 12 cattle notifications were confirmed BTV-3/NET2023 positive as confirmed by PCR-testing. In the second week (week 37) the total number of cases increased to 18 in sheep flocks and 55 in cattle herds. In the third week (week 38), the total number of cases increased to 324 in sheep flocks and 61 in cattle herds. An geographical overview of the potential index cases and the spread of BTV-confirmed suspicions is shown in **Figure 4**.

**Figure 4.**
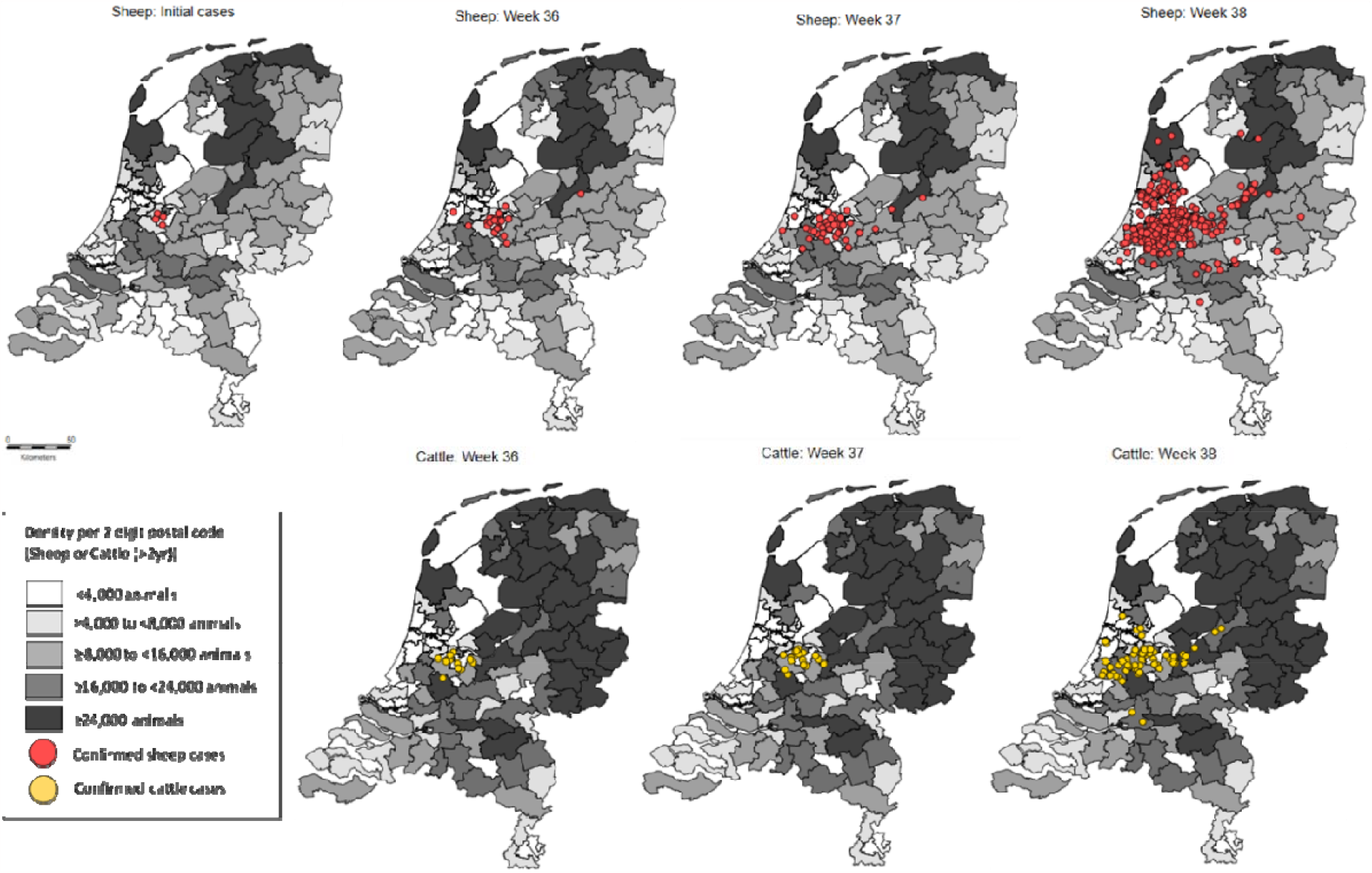
Thematic map of the Netherlands with the number of sheep or cattle density per 2 digit postal code. BTV-confirmed cases of sheep (red dots) and cattle (yellow) of the first (week 36), second (week 37) and third (week 38) week of the bluetongue serotype 3 outbreak are also depicted.

### Retrospective study

To investigate whether the initial outbreak started at the four farms in the middle of the Netherlands, bulk tank milk samples from cattle farms submitted for routine testing in August were screened for BTV-antibodies. Antibody positive results were found in 2.8% (95% CI: 1.9-4.1%) of the bulk tank milk samples. Of the 991 bulk tank milk samples, 955 tested negative, eight tested doubtful and 28 tested positive. However, 24 out of 36 bulk tank milk samples with doubtful or positive results (58%) were from farms with a proven history of vaccination against BTV-8. BTV antibodies that could not be linked to a recorded history of vaccination against BT were found in 12 out of 991 bulk tank milk samples. These 12 positive samples were however not clustered and showed a somewhat similar distribution as the positive samples that originated from vaccinated herds **(Figure 5)**. Altogether, these results show that there was no area with a high seroprevalence for BTV antibodies in August 2023 and no BTV-specific antibodies were found in the region were the initial notifications of BTV were made. Therefore, the four index cases where bluetongue was initially observed belong more likely to one of the first affected farms.

**Figure 5.**
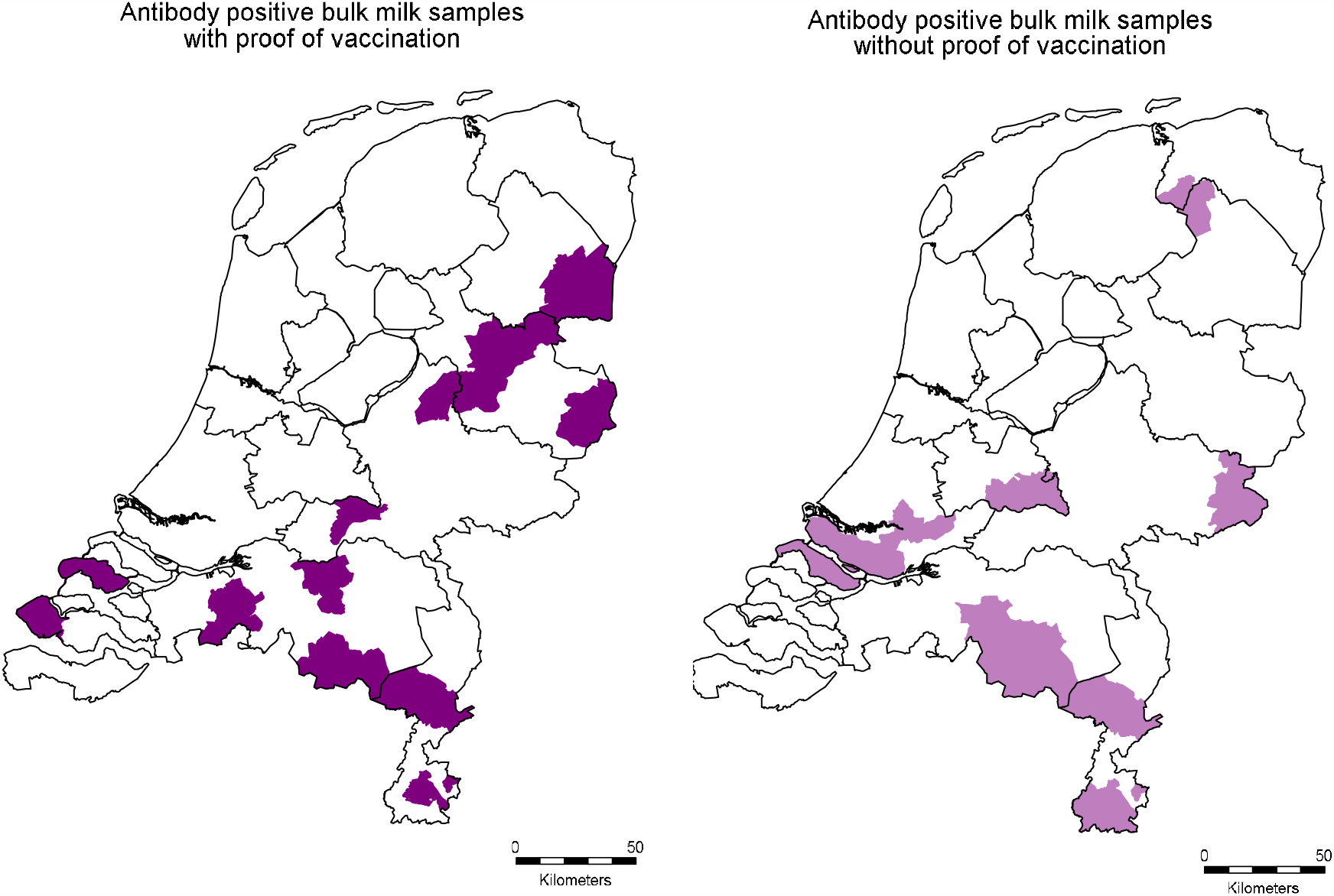
Distribution of location from which the bulk milk samples originated that tested doubtful or positive for antibodies against BTV, for herds where <u>evidence</u> of vaccination in the last five years was found (left) or <u>no evidence</u> of vaccination in the most recent five years was found (right)

## Discussion

This manuscript describes the actions after the emergence of a novel BTV-3 strain in sheep and cattle in the Netherlands. Initially, sheep from four farms located in the middle of the Netherlands showed clinical signs of fever, lethargy, hypersalivation, ulcerations, erosions of the oral and nasal mucosal membranes or sudden death. These sheep were tested positive by real-time PCR and all but one showed seroconversion by competition ELISA. That one sheep was acutely infected and was still negative for BTV antibodies. Whole genome sequencing using Oxford Nanopore Technology has shown that the full viral genome sequence can be determined quickly. The generated nucleotide sequence of segment 2 aligns with other sequences from BTV serotype 3. We investigated the epidemiological situation the month before the first cases were found by retrospectively testing of bulk tank milk, however, no high seroprevalence was measured nor seropositive samples were found in the region were the initial cases were detected, indicating that the index cases of the initial four farms could be considered as one of the first BTV infections. This was in agreement with the findings of retrospective analysis of 1003 sheep sera from 89 flocks which indicated 3,4% herd prevalence in August (data not shown). Currently, on 29 September 2023, 380 farms or holdings have been confirmed positive by real-time PCR.

Early detection of diseases by clinical diagnosis remains challenging, especially for unpredicted non-endemic diseases. Like for bluetongue, the condition knows a wide and non-specific spectrum of clinical manifestations as fever, hypersalivation, lameness, edemas and sudden death. The variation in disease severity of bluetongue in sheep and cattle overlaps with a number of endemic infections. For instance, diseases like orf, dermatophilosis, haemonchosis, pasteurellosis, (strawberry) footrot and photosensitisation are differentially diagnostic relevant endemic conditions in sheep. Malignant catarrhal fever and photosensitisation can cause similar signs as BT in cattle (24,25). Awareness of BTV-like symptoms among veterinarians is also of great importance for other notifiable diseases like FMD, peste des petits ruminants (PPR), sheep and goat pox (SGP), and emerging haemorrhagic disease (EHD) and should, in case of suspicion, be notified to the official authorities. During an outbreak of BTV, publicity creates increased awareness among veterinarians and farmers. This might result in an increase of false notifications of bluetongue based on the, not so specific, clinical signs. Therefore, education of veterinarians and livestock farmers about the clinical manifestations of BT and other diseases remains important. In particular of notifiable diseases of which countries are free of for a longer time since many veterinarians may have not seen the clinical picture in real life. Only laboratory diagnosis can and should rapidly differentiate between these notifiable diseases to support clinical diagnosis. Nevertheless, in this outbreak, this emerging disease has been detected successfully in an early stage.

The speed of the onward spread of BTV after the initial emergence clearly shows that indigenous *Culicoides* in the Netherlands are vector competent to transmit the causative BTV-3/NET2023. Since the predominantly Afro-Asiatic vector of BTV, C. *imicola*, is not present in Northwestern Europe, and the BTV-6 outbreak in the Netherlands in 2008 showed that the outbreak dies out when indigenous *Culicoides* are unable to effectively transmit the virus, BTV-3/NET2023 is after BTV-8/NET2006 the second BTV variant successfully transmitted by Northwestern indigenous midge species (26). Previously, it has been shown that BTV-8/NET2006 is transmitted by indigenous midge species of the *Culicoides obsoletus* complex, such as C. *obsoletus*, C. *scoticus*, C. *dewulfi* and C. *chiopterus* (27–29). Entomological research is needed to identify the midge species involved in transmission of BTV-3/NET2023.

Immediate arising questions are the geographical origin and route of introduction of BTV-3 into the Netherlands. Phylogenetic analysis of Seg-2 clearly shows clustering with other Seg-2 sequences of serotype 3, including geographically close variants of BTV-3. However, sequences of other genome segments do not show any undoubtful high homology with one of the BTV-3 variants. Therefore, tracing back where the virus geographically is coming from is difficult to determine. Furthermore, the segmented genome of BTV is the basis for exchange of genome segments known as reassortment, including antigenic shift. This viral trait hampers the unravelling of the geographical source of BTV-3/NET2023. It can be speculated that the virus was introduced over a long distance as neighboring countries Belgium and Germany recently obtained the BTV-free status since June 2023. In conclusion, no clue can be given upon the source and route of introduction of BTV-3/NET2023. However, yearly monitoring, and the in this paper presented retrospective study showed that virus circulation only recently started in the currently affected area.

In conclusion, after a decade of BT-freedom, BTV-3 emerged in the Netherlands causing clinical signs and mortality in sheep and cattle. The causative virus is designated BTV-3/NET2023. The source, geographical origin, and introduction route of BTV-3/NET2023 are unknown but virus circulation has recently started in the currently affected area. Clearly, BTV-3/NET2023 is transmitted by indigenous Dutch midges but the involved midge species is/are not identified yet.

## Acknowledgements

We would like to thank the veterinary practitioners from Dierenartsenpraktijk Gorter and Dierenkliniek Amstel, Vecht en Venen. Additionally, we want to thank all the laboratory personnel at WBVR and Royal GD and all veterinarians that conducted the herd visits for their laborious work. This study is financially supported by the Ministry of Agriculture, Nature and Food quality with projectnumber: 1600002757 VZVD.

## Biosketch

Melle Holwerda is head of the national reference laboratory for viral veterinary vector-borne diseases at WBVR. Inge Santman-Berends works as head of the department of epidemiology at the Royal GD.

